# The complex optic lobe of dragonflies

**DOI:** 10.1101/2020.05.10.087437

**Authors:** Joseph M. Fabian, Basil el Jundi, Steven D. Wiederman, David C. O’Carroll

## Abstract

Dragonflies represent an ancient lineage of visual predators, which last shared a common ancestor with insect groups such as dipteran flies in the early Devonian, 406 million years ago [1,2]. Despite their important evolutionary status, and recent interest in them as a model for complex visual physiology and behavior, the most recent detailed description of the dragonfly optic lobe is itself more than a century old [3]. Many insects process visual information in optic lobes comprising 4 sequential, retinotopically organized neuropils: the lamina, medulla, lobula and a posterior lobula plate devoted to processing information about wide-field motion stimuli [4, 5]. Recent reports suggest that the dragonflies also follow this basic plan, with a divided lobula similar to those of flies, moths and butterflies [6, 7]. Here we refute this claim, showing that dragonflies have an unprecedentedly complex lobula comprising at least 4 sequential synaptic neuropils, in addition to two lobula plate like structures located on opposite sides of the brain. The second and third optic ganglia contain approximately twice as many synaptic layers as any other insect group yet studied. Using intracellular recording and labeling of neurons we further show that the most anterior lobe contains wide-field motion processing tangential neurons similar to those of the posterior lobula plate of dipteran flies. In addition to describing what is probably the most complex and unique optic lobe of any insect to date, our findings provide interesting insights to understanding the evolution of the insect optic lobe and serve as a reminder that the highly studied visual circuits of dipteran flies represent just a single derived form of these brain structures.

## Results and Discussion

### The dragonfly optic lobe

The major divisions of synaptic neuropil in the dragonfly optic lobe (*Hemicordulia tau*) are clearly evident from confocal images of thick horizontal sections labelled using antibodies against the presynaptic vesicle protein synorf1 (Fig. 1A). Here fluorescence corresponds to regions of high synaptic density within neuropil, while darker areas represent dividing layers of cell bodies or the major axon tracts that serially interlink neuropils [8]. On this basis, the large and complex optic lobes can be readily segmented into 11 distinct neuropils; the lamina, the outer medulla, serpentine medulla, inner medulla, an anterior accessory medulla (not visible in Fig. 1A), and a lobula complex with 6 distinct subdivisions (Fig. 1A,B). We observed this same basic plan in synapsin stained sections and whole-mount brains of species from 3 families: the Corduliidae (Fig. 1, S1), Libellulidae, and the Aeshnidae (Fig. 2). We segmented confocal image stacks from wholemount preparations in three-dimensions (3D) to produce digital reconstructions of identifiable neuropils (lamina removed during wholemount preparation) in individual brains across all three species (Fig. 2B). In relative terms, the dragonfly medulla is extremely large, as is expected given the compound eye holds up to 24000 facets [9]. Although absolute brain size differs across this group (correlated with obvious large differences in body size) the optic lobe organisation observed in *Hemicordulia* (Fig. 1) is conserved across all three families. These families last shared a common ancestor in the early Triassic, some 250 million years ago [2], with the Aeshnidae representing the most ancient lineage of extant dragonflies, so this complex lobula organisation is likely plesiomorphic in odonates.

**Figure 1:**
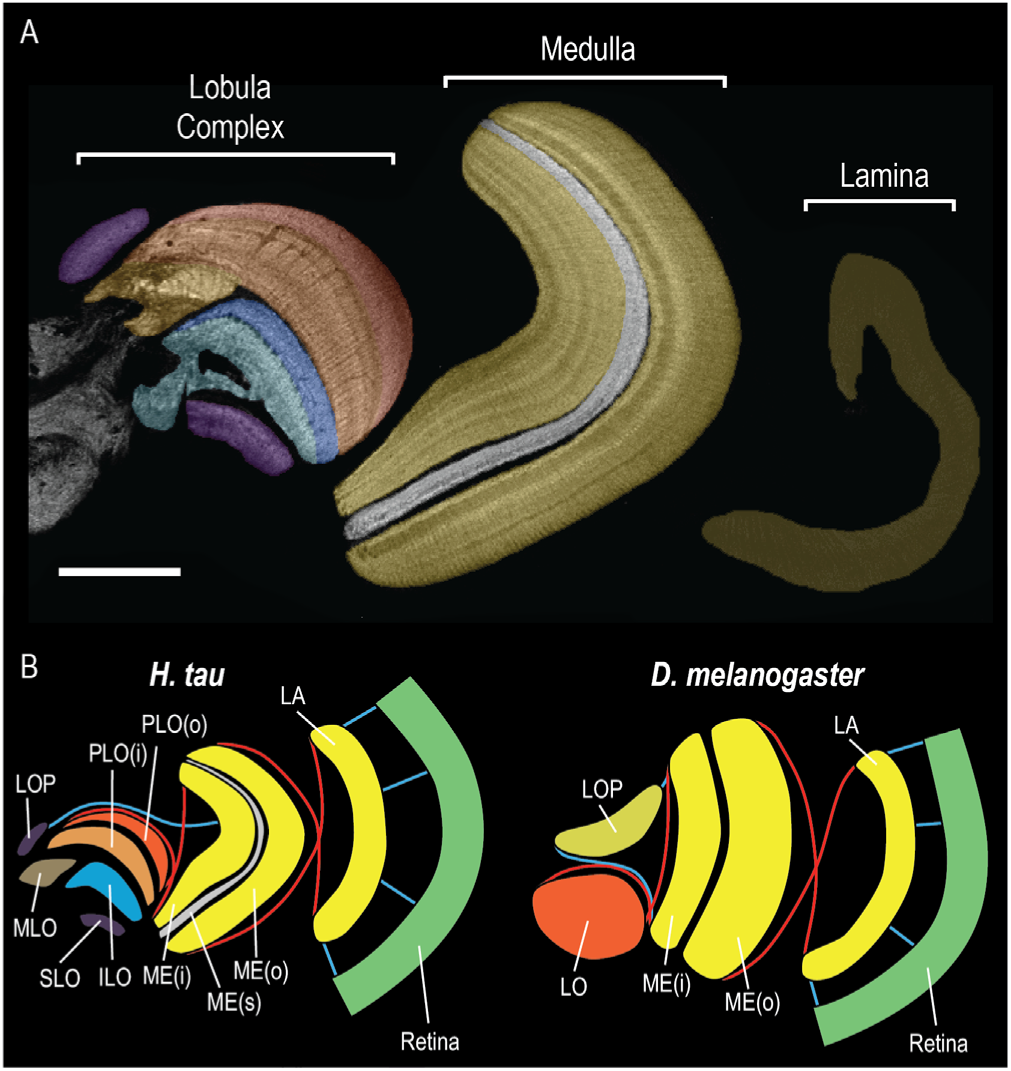
The complex optic lobe of *Hemicordulia tau*. (A) A synapsin stained horizontal section through the optic lobe of *H. tau*, defining the segmentation of different neuropils based on anti-synapsin immunoreactivity. Scale bar = 200 μm. (B) An diagrammatic view of the optic lobe of *H. tau* (left) and *Drosophila melanogaster* (right). LA = Lamina, ME(o), ME(s) and ME(i) = outer, serpentine and inner medulla, PLO(o) and PLO(i) = outer and inner subunit of the Primary Lobula, ILO = Inner lobula, SLO = Sublobula, MLO = Medial Lobula and LOP = Lobula Plate. Blue lines indicate major uncrossed serial connections, while red lines indicate crossed connections (chiasmata), see also Figure S1.

**Figure 2.**
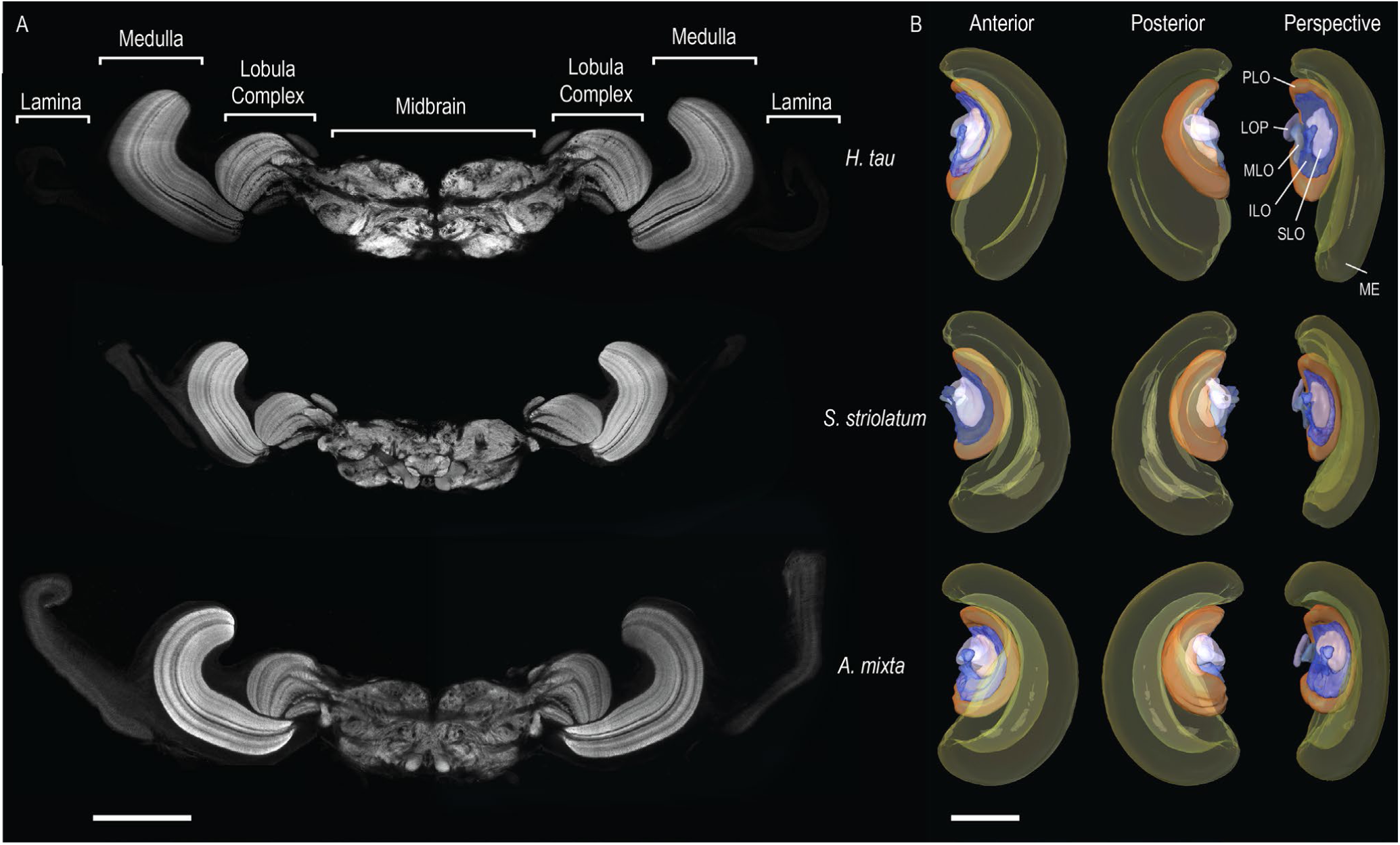
Similarity of the optic lobes in three dragonfly species. (A) Synapsin stained horizontal sections through the brain of dragonflies. Each section displays the left and right optic lobes, flanking the central protocerebrum. Depths within the optic lobe are similar across sections, however the corresponding protocerebral depth varies. (B) 3-dimensional reconstructions of the optic lobe across all three species. Reconstructions were produced by segmenting identified structures from confocal images of individual wholemount brains stained for synapsin. Structures were labelled in accordance with figure 1B, however the inner and outer subunits of the medulla and primary lobula are reconstructed as a single structure for clarity. Scale bars = 500μm.

The organisation of the dragonfly lobula is clearly more elaborate than those of holometabolous insects including dipteran flies such as *Drosophila melanogaster* [6,10](Fig 1B), Coleoptera [11,12] and Lepidoptera [13-17]. In these insects, the lobula is divided into two subunits, a posterior lobula plate that receives direct projections from the inner medulla (ME(i)) and a larger anterior lobula that receives inputs that cross over at a chiasm with the inner medulla (ME(i), fig. 1B). In the dragonflies, this prominent chiasm is also present between the medulla and the outermost and largest of a series of at least 4 neighbouring lobula neuropils. The primary lobula (PLO, fig 1B), spanning the entire distal surface of the lobula complex is divided into striate outer (PLO(o)) and inner (PLO(i)) regions by a prominent dark serpentine layer very similar to those seen in the medulla of many insects [18]. Nested within the proximal surface of the PLO(i) there are then two smaller structures that we term the medial and inner lobula (MLO, ILO, fig. 1B). The wedge-shaped MLO intrudes between the posterior portions of the primary and the inner lobula. The inner lobula lies anterior to this medial unit, and is further divided into outer and inner subunits by a thin layer of lower synaptic density, with the innermost part extending further medially into additional synaptic structures which may be contiguous with those of the lateral mid-brain (Fig. 1A). The columnar appearance of both the MLO and ILO and the absence of obvious additional tracts linking these structures directly to medulla projections suggests that they are postsynaptic to the primary lobula and retinotopically organised, thus forming two additional 4^th^ optic ganglia.

Moreover, lying anterior and medial to the inner lobula, is a further distinctive structure that we term the sublobula (SLO, Fig 1 A,B), by analogy to a term originally proposed by Cajal and Sanchez [4] for the inner part of the lobula in honeybees. Strausfeld [7] suggests that the sublobula of bees may be functionally equivalent to the dipteran lobula plate as a site of wide-field motion processing. In the honeybee it is fused to the outer lobula as a single, large, synaptically dense structure [19,20], but in the dragonfly this structure is clearly segregated from the inner lobula by a thick band devoid of synapsin immunoreactivity. Finally, a small 6^th^ subunit sits on the posterior surface of the lobula complex, at the medial margin of the outer lobula. Osmium stained horizontal sections and injections of neuronal tracer into the posterior medulla (Fig. S1) reveal direct projections of a population of columnar neurons into this subunit, posterior to the second optic chiasm (OCH2, Fig S1), so it likely lies at a similar level of hierarchical processing to the outer primary lobula (PLO(o)) and the lobula plate of holometabolous insects [6,7]. Outside the holometabola, relatively small satellite neuropils that receive uncrossed axons and lying to one side or beneath the lobula have also been observed in Mantoidae and plecopterans (stone flies) [7].

### Layering of the Medulla and Lobula complex

One of the most striking differences between the dragonfly optic lobes and those of other insect species is in the extreme degree of concentric stratification seen in each of the major divisions of the optic lobes. We further investigated the complexity of individual neuropils by identifying individual synaptic layers and attempting to compare these with the layers identified in the medulla and lobula of other insect species. This is complicated by the fact that recent work has numbered layers of different species based on inconsistent criteria [15,21,22]. Perhaps the most robust numbering of layers comes from classical studies using Golgi and other techniques to identify 10 key strata in the fly medulla based on apparent common synaptic layers of key classes of columnar and tangential neurons [18,23]. In an attempt to provide a more robust basis for identifying layers than used in prior work, we computed pixel intensity plots from confocal images of sectioned *H. tau* brains co-labelled with antibodies against synapsin and serotonin. These two labels are complementary since synapsin is widely expressed in areas of high synaptic density [8], allowing visualisation of key input and output layers, while extensive anti-serotonin immunoreactivity has been observed in large tangential cells that arborize across the medulla of insects [24] providing a general stain that selectively highlights tangential neuronal structures, greatly improving layer discriminability (see fig S2 for a complete section). To minimise the possibility of subjective errors we generated six maximum intensity projections of local 7.5 μm confocal stacks at different depths from thick horizontal sections through the medulla (Fig 3 A-C) and primary lobula (Fig 3 D-E). Synaptic layers were defined by contrast boundaries in either anti-synapsin or anti-serotonin immunoreactivity observed across multiple depths in the sample.

**Figure 3.**
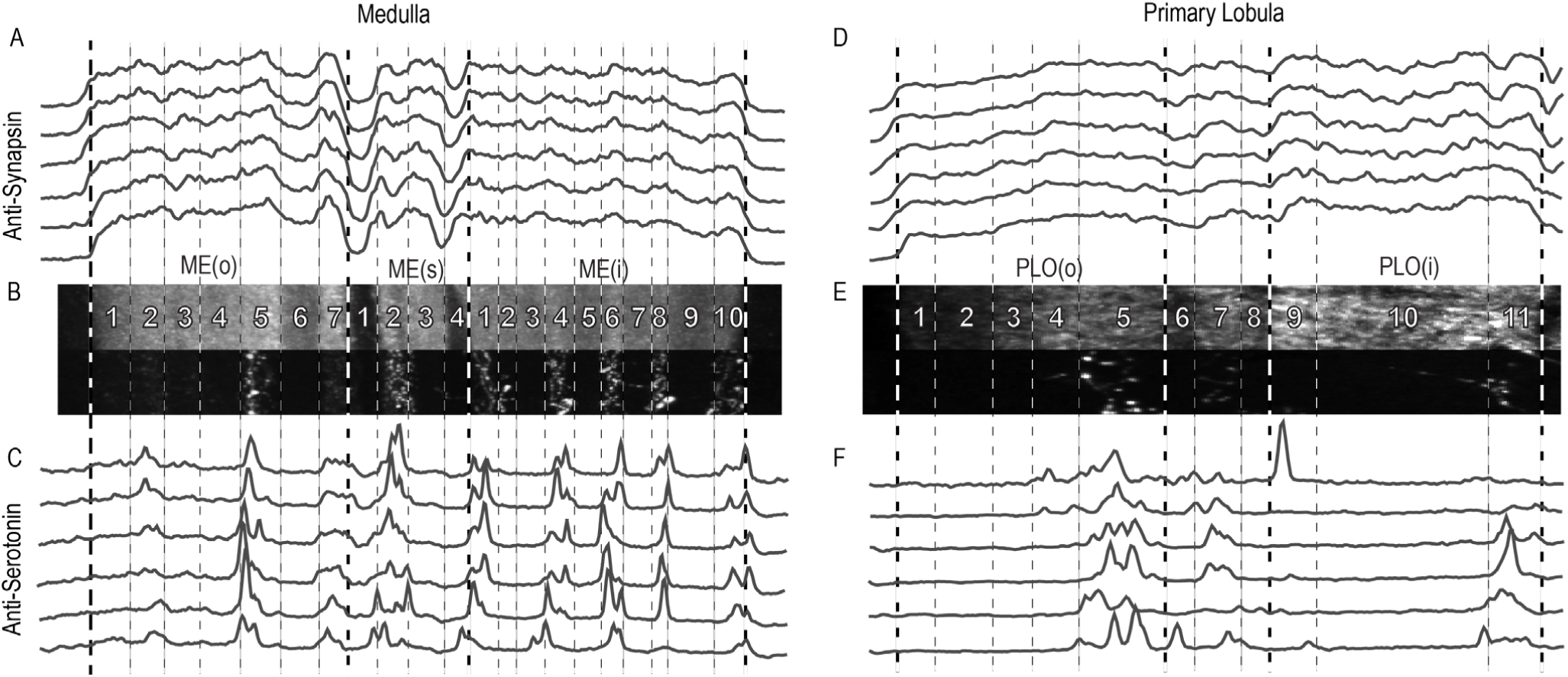
The synaptic layers of the medulla and lobula in *Hemicordulia*. (A) Synapsin immunoreactivity intensity plots from a series of maximum intensity projections through the medulla obtained from different depths of a confocal image stack from a horizontal section obtained from *Hemicordulia*. (B, E) Example confocal images of anti-synapsin (top) and anti-serotonin (bottom) immunoreactivity, used to generate intensity plots. (C) Serotonin immunoreactivity intensity plots from the corresponding positions in A. (D) The synapsin immunoreactivity of the lobula. (F) Serotonin immunoreactivity intensity plots from the corresponding positions in D.

Based on this analysis, we can distinguish 21 layers in the dragonfly medulla, compared to the 8-11 layers observed in all other insects studied [15,21-23, 25]. Given this unprecedented complexity, specific comparisons between individual layers and those of other insects is difficult. For example, the 9^th^ and 10^th^ medulla layers lie within a mid-band of synapsin staining between 2 dark bands that may be homologous with the dipteran serpentine layer (M7), but we then see a highly complex subdivision of 10 additional layers in the inner medulla (compared with 2-3 in Diptera). Therefore, while we adopt the naming conventions recently recommended for the major divisions of these ganglia [6], we implement a separate numbering system for the layers of the outer medulla, serpentine region and inner medulla.

We identify a total of 7 layers (MEO1-MEO7) in the outer medulla, 10 in the inner medulla (MEI1-Mi10) and 4 in a ‘serpentine’ region (S1-S4) that separates the inner from the outer medulla. This region is bounded by dark layers S1 and S4 which show little, if any synapsin staining, and likely contain the axons and major dendrites of tangential and amacrine processes that feed into the medulla in many insects [18]. Whilst these layers are not technically ‘synaptic’, we include them in our analysis to be consistent with other studies where a single serpentine layer is commonly reported as medulla layer M7 [15,18]. The function of the dragonfly dual serpentine layer and the two synaptic layers within it, are currently unknown, although anti-serotonin immunoreactivity is especially prominent in layer S2. Indeed, of the 21 layers in the dragonfly medulla, 9 show consistent anti-serotonin immunoreactivity (Fig 3).

Whilst the absolute number of layers is striking, their distribution is equally unusual. In most insects the outer medulla houses 6 synaptic layers, compared to just 2-3 in the inner medulla. Our findings show that the synaptic organisation in the dragonfly medulla is the opposite, with an extremely elaborate inner subunit consisting of 10 layers - the same as the total number of layers observed in the *Drosophila* medulla. No studies have yet investigated the physiological function or cell types present in these layers of dragonflies, but in other species these are the major input sites for retinotopic columnar neurons that project to the lobula complex. The large number of such layers in the dragonfly may prove to be linked to the larger number of possible post-synaptic targets in the highly divided inner layers of the lobula complex.

The highly layered organisation evident from the medulla is repeated within the lobula. Within the primary lobula alone we observed 11 layers (L1-L11). As with layers S1 and S4 of the medulla, Primary Lobula layers L6 and L8 are serpentine-like ‘dark’ layers, with weak anti-synapsin immunoreactivity that separates the inner and outer Primary Lobula. Indeed the shape, size and number of synaptic layers in the primary lobula is so exaggerated that it resembles the medulla of many other insect species. While there has not yet been any complete description of all the cell types present, the outer layers of the Primary lobula host the dendrites of several previously described target-detecting neurons [26,27]. While we did not quantitatively analyse layering of the higher-order lobula subunits, the inner lobula (Fig. 1) and sublobula also show a serpentine-like dark mid-bands so could potentially be segregated further into outer and inner divisions. Hence the total number of lobula layers is certainly more than the 11 of the primary lobula.

### The sub-lobula and wide field motion analysis

In considering the substantial differences between the dragonfly lobula complex and that of other insects, one interesting question that emerges is that of which subunit serves the role served by the lobula plate of other species. To date, most research on the lobula plate of species such as dipterans supports it having a primary role in visual motion analysis, while other visual tasks (e.g. polarization vision, pattern analysis, colour processing or feature detection) may be handled by neural circuits of the lobula and mid-brain [28-31]. The widely studied Lobula Plate Tangential Cells (LPTCs) are an important group of motion-sensitive neurons found in a variety of flying insect species [32] and which mediate responses to optic-flow cues. This is a visual sub-modality that dragonflies would also encounter during their spectacular hovering flight and acrobatic aerial pursuits of conspecifics and prey, which in many respects rival those of territorial Diptera with advanced visual abilities. However, despite their ancient lineage - among the oldest of the extant insect groups – the most obvious structure that we identified as a posterior lobula plate in the dragonfly (Fig. 1, S1) is approximately 12 fold smaller in volume terms (Table S1) than the lobula plate of other insects with similar sized eyes and brain, such as the male sphingid moth *Manduca* [14] (e.g. total brain volume, excluding the lamina = 6.81×10^8^ μm^3^ in *Hemicordulia* versus 6.95 × 10^8^ μm^3^ in *Manduca*).

We recently identified a diverse population of neurons in the dragonfly with similar physiological characteristics to LPTCs of other insects [33]. To investigate where these dragonfly lobula tangential cells (LTCs) lie in the dragonfly lobula complex, we combined intracellular recording techniques with fluorescent dye labelling (fig. 4). Fig. 4A shows the responses of one such neuron to stimulation with a sinusoidal grating. Motion in the preferred direction elicits a robust depolarisation and increase in spiking activity, whilst gratings moving in the opposite direction produce hyperpolarisation. These responses strongly resemble those seen in direction-selective HS or VS neurons of the fly lobula plate [32]. Wholemount confocal images of intracellularly labelled neurons reveal broad spiny input dendrites that span the medial and anterior portion of the lobula complex (fig 4B, D). Horizontal sections from these same brains were subsequently immunolabelled with anti-synapsin antibodies to reveal the synaptic sites of these inputs. This reveals that both neurons receive inputs in the sublobula (fig 4C, E).

**Figure 4.**
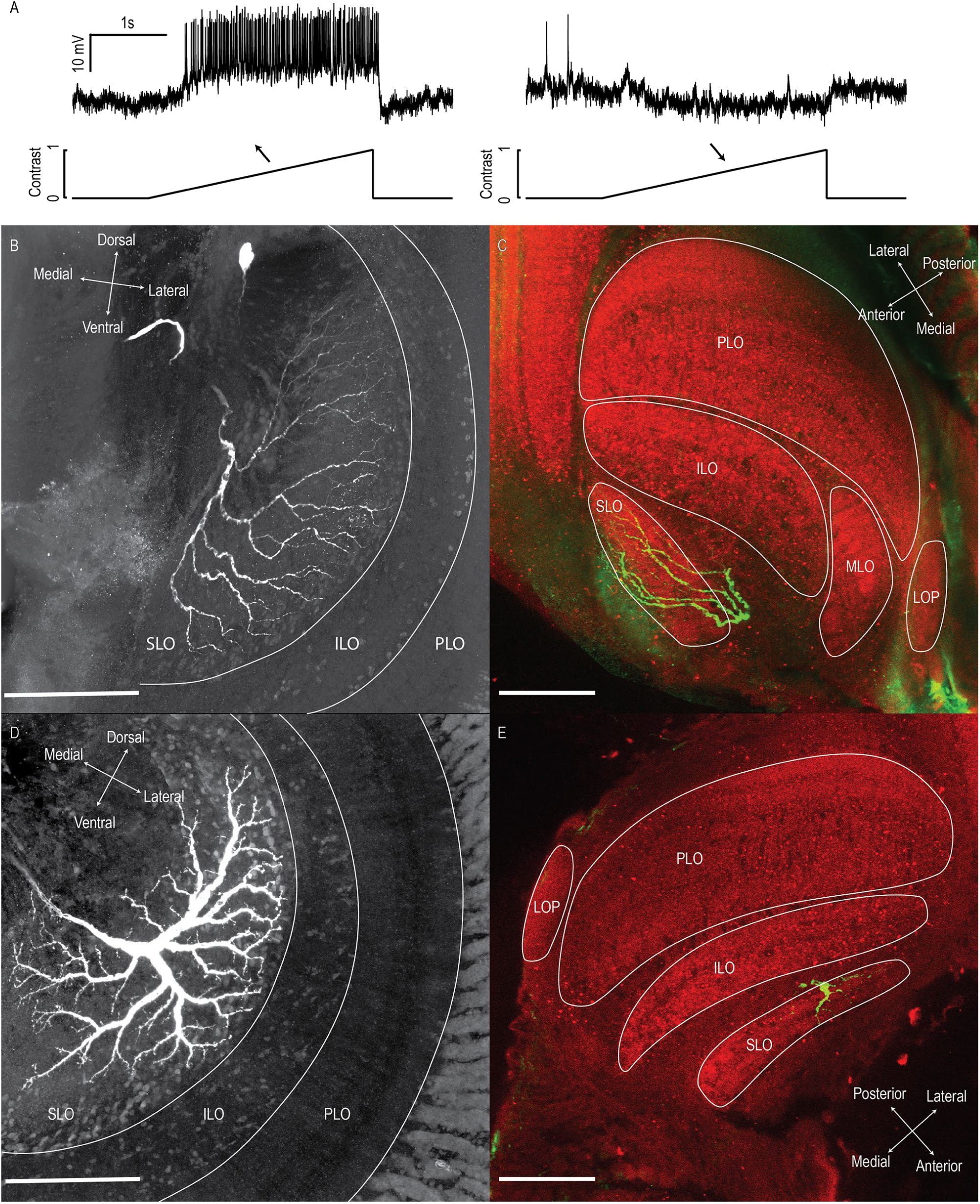
Motion-sensitive neurons of the dragonfly sublobula. (A) A subset of neurons in the dragonfly lobula complex provide robust, direction opponent responses to drifting gratings. (B) A wholemount montage of a dye-filled bar-sensitive neuron labelled by intracellular lucifer yellow injection. (B) Horizontal Sections through the lobula complex confirm the input dendrites of this motion sensitive neuron lies within the Sublobula. (C) A wholemount montage of an optic flow sensitive neuron. (D) Neurons sensitive to optic flow also receive inputs in the sublobula.

### Evolution of the divided lobula

Our results conclusively demonstrate that the dragonfly sublobula most certainly contains a population of tangential neurons closely resembling those of LPTCs in dipteran flies. It is therefore difficult to reconcile our findings with recent neuroanatomical studies [7] and a review of the invertebrate brain [6] that grouped dragonflies along with holometabolous Diptera, Lepidoptera and Coleoptera as having a divided lobula with a lobula plate. We did indeed find a small lobula plate-like structure on the posterior surface of the optic lobes. Prior work with reduced silver preparations reveals neurons with both columnar and tangential processes within a small posterior lobula plate [7] and our own confocal imaging of anti-synapsin stained material also reveals dark shadows of tangential processes in this region. Since we have yet to record from tangential neurons that arborise exclusively in this region, we cannot preclude the possibility of a second sub-population of tangential neurons displaying similar properties to those we have found in the sublobula. Nevertheless, our results suggest that key functions of the lobula plate of more recent insect groups is handled by a separate structure, on the other side of the brain.

Despite dragonflies representing an ancient lineage, it is of course possible that their highly complex lobula arrangement may be a highly derived trait, specialised to the unique needs of the apex predator of the flying insect world. Nevertheless, an alternative possibility is that it is the simpler divided lobula of holometabola that is the derived state. Indeed, a motion sensitive sublobula structure is not unique to dragonflies, with wide-field motion sensitive neurons also described from a similar region of the honeybee lobula [34,35]. While Strausfeld [7] previously argued that the sublobula of bees may be supplied by homologues of the medulla T4 neurons that feed the lobula plate in other holometabolous insects, this view of the sublobula as a homologue of the dipteran lobula plate seems at odds with our finding that the dragonfly has both structures, on opposite sides of the brain. Dragonfly equivalents of the dipteran medulla T4 and lobula T5 neurons have yet to be identified. Since these can be considered to be sibling cells that might have arisen by the duplication of progenitors in a common ancestor [36], further analysis of the columnar neurons that project from the proximal medulla to different lobula subunits in the dragonfly may shed further light on whether the sublobula is indeed a homologue of the dipteran lobula plate.

## Materials and Methods

### Experimental Model and Subject Details

#### Dragonflies

Male dragonflies were caught in the wild, either from the Adelaide Botanical Gardens, Australia (*Hemicordulia tau, Adversaechna Brevistyla*) or various wetlands in Lund, Sweden (*Aeshna mixta, Sympetrum striolatum*). Animals were stored in small, moist plastic bags at 4°C until use, normally not exceeding 5 hours.

#### Wholemounts

To visualise neuropil in wholemount specimens, we followed the staining protocol by Ott [37]. We carefully dissected and removed the brain from the head capsule under HEPES buffered saline (HBS; 150 mM NaCl; 5 mM KCl; 5 mM CaCl2; 25 mM sucrose; 10 mM HEPES; pH 7.4) before fixation for 24 hours at room temperature (RT) in zinc-formaldehyde solution (ZnFA; 0.25% [18.4 mM] ZnCl2; 0.788% [135 mM] NaCl; 1.2% [35 mM] sucrose; 1% formaldehyde). Following fixation, brains were rinsed in HBS (8 × 20 minutes), before treatment in 80/20 DMSO/methanol solution for 55 minutes at RT. Brains were then rinsed in Tris-HCL (3×10 minutes) and preincubated in 5% Normal Goat Serum (NGS) for 3 hours at RT or overnight at 4°C. Primary antibodies (anti-synapsin, RRID:AB_528479) were diluted 1:50 in PBT and 1% NGS, and incubated for 5 days at 4°C under gentle agitation. Brains were then rinsed (8×20 minutes) in PBT, and treated with secondary antibodies (Goat anti-mouse CY5, RRID:AB_10895546) diluted 1:300 in PBT and 1% NGS for 3 days at 4°C under gentle agitation. Brains were then rinsed (6×20 minutes) in PBS, and dehydrated by ascending ethanol series (50%, 70%, 80%, 90%, 100%, 15 minutes each). Dehydrated brains were cleared in Methyl salicylate, (50% Methyl salicylate 50% ethanol for 15 minutes, 100% Methyl salicylate for 1 hour). Finally brains were mounted in Permount between two #1.5 coverslips, separated by plastic adhesive spacer rings (∼500μm).

#### Sections

Sections for synapsin staining followed the exact same protocol as above, with the following exceptions: Following primary fixation, brains were embedded in a gelatin-albumin mixture (4.8% gelatin and 12% ovalbumin in milliQ water), and allowed to set at RT before post-fixing overnight in 4% Paraformaldehyde at 4°C. 200μm horizontal sections were cut on a vibratome before rinsing (6×20 minutes) in PBS. No DMSO step was included due to the reduced thickness of tissue. Sections were either incubated with anti-synapsin alone, or in combination with anti-serotonin or anti-lucifer yellow (*H. tau* only). Primary antibody incubation was reduced to 3 days. For sections that included intracellularly labelled neurons, synapsin was visualised with a Goat anti-mouse CY3 secondary antibody. For sections co-labelled for serotonin, this was visualised with a Goat anti-rabbit Alexa Fluor 546 secondary antibody.

Additional thick sections for osmium staining were prepared from from one individual of *Hemicordulia* from a brain fixed in 2% glutaraldehyde, 4% paraformaldehyde in 0.2 M phosphate buffer (+5% sucrose) at pH 7.4 for 2 h. Tissue was rinsed in PBS (2×10 minutes) before embedding in 4% agarose (in buffer), allowed to cool to just above setting point (ca. 25°C). After cooling, the block was cut into 100 μm sections on a vibratome, which were then post-fixed in 1% aqueous OsO4 for 30 min before dehydration through an ethanol series, and mounting under coverslips in unpolymerized araldite.

#### Electrophysiology and intracellular neuron labelling

*Hemicordulia tau* dragonflies were immobilised with a 1:1 wax/rosin mixture and fixed to an articulating magnetic stand. A small hole was dissected on the posterior surface of the head capsule directly above the left lobula complex. Aluminosilicate electrodes were pulled on a Sutter Instruments P-97 electrode puller, and tip filled with 4% Lucifer Yellow in 1M LiCl solution. Electrodes had typical resistances of 150-200 MΩ. Electrodes were placed in the medial portion of the lobula complex, and stepped through the brain from the posterior to anterior lobula complex using a piezo-electric stepper (Marzhauser-Wetzlar PM-10). Intracellular responses were digitized at 5 kHz with a 16-bit A/D converter (National Instruments) for off-line analysis with MATLAB. Following physiological characterisation, neurons were injected with approximately -2nA of current for up to 30 minutes. Immediately following injection the brain was carefully dissected under PBS, before fixation overnight in 4% paraformaldehyde at 4°C. Brains were then processed in accordance with published protocols [38]. Brains were rinsed (3×10 minutes) in PBS, before permeabilization in 80/20 DMSO/Methanol solution for 55 minutes. Brains were then rinsed (3×30 minutes) in PBT, and preincubated in 5% NGS in PBT for 3 hours at RT under gentle agitation. Brains were incubated in 1:50 dilution of primary antibody (anti-lucifer yellow, RRID:AB_2536190) in universal antibody dilution solution (Sigma Aldrich) for 3 days at 4°C under gentle agitation. They were then rinsed (3×30 minutes) in 10% NGS, before incubation with a 1:50 dilution of NeutraAvadin DyLight 633 for 3 days at 4°C under gentle agitation. The samples were then rinsed, dehydrated and mounted identically to wholemount synapsin labelled samples described above. Following confocal imaging brains were retrieved from coverslips with xylene, rehydrated by an inverted ethanol series (90%, 80%, 70%, 50%, 0%), and sectioned through an identical protocol as described above.

#### Visual Stimuli and Physiological Characterisation

Recordings obtained were during ongoing experiments aimed at classifying motion sensitive and feature selective neurons such as the small target motion detecting (STMD) neurons and lobula tangential cells (LTC’s) described in our recent work [33, 39-41], from more than 300 dragonflies over a 4 year period. Upon establishing a healthy recording, all neurons were characterized using a range of stimuli presented on an LCD monitor (either an Eizo Foris FG2421 LCD at a frame rate of 120 Hz, or an Asus ROG Swift PG279Q IPS LCD at 165 Hz). Stimuli included a sequence of drifting texture patterns, drifting sinusoidal gratings with varying directions, spatial and temporal frequency, and discrete features (e.g. targets, bars) of different sizes and contrast. In addition to hundreds of recorded neurons that were primarily selective for discrete features such as small targets and which gave weak responses to optic flow stimuli, we obtained reasonably complete data sets for 56 lobula neurons that provided robust responses to wide-field motion stimuli. Despite all giving strong responses to optical flow stimuli, these neurons nevertheless display very diverse physiological properties, with some displaying non-directional responses or strong responses to discrete edges or features, so ongoing work is required to establish a classification of their basic properties. One subset, however, gave robust, direction opponent responses to wide-field texture patterns and gratings, with little to no response to small moving features. Most of this subset of neurons also show the sensitivity to contrast and the tuning and selectivity for spatial and temporal frequency very similar to neurons classified as lobula plate tangential cells (LPTCs) in dipteran flies and other taxa [32].

#### Tracer injection

*Aeshna mixta* dragonflies were immobilised with a 1:1 wax/rosin mixture and fixed to an articulating magnetic stand. A large hole was cut in the posterior head capsule, allowing visualisation of the optic lobe. The neural sheath was carefully removed, and any excess liquid was absorbed with tissue paper. Aluminosilicate microelectrodes had their tips manually broken and were then dipped in Vaseline, before dipping into fluorescent dextran 488 (3000 mw.) crystals. The electrode was then inserted into the posterior part of the left medulla by hand. After thoroughly rinsing the head capsule with PBS to ensure that no crystals remained on the brain surface, animals were placed in a small moist container, and left overnight at 4°C. The following day, the brain was carefully dissected under PBS and fixed overnight in 4% PFA. Brains were then washed, dehydrated, cleared and mounted as described previously.

#### Imaging, Image Processing and 3D Reconstruction

Samples were imaged on a confocal microscope (Zeiss LSM 510 Meta), using a 10x (Zeiss C-Apochromat, NA=0.45 water immersion) or 25× (Zeiss LD LCI Plan-Apochromat, NA=0.8, with correction for oil immersion) objectives. Samples were scanned using either a 561 (Cy3) or 633 (Cy5, Neutravadin Dylight 633) laser line. All samples were imaged as a Z-series at a resolution of either 1024×1024 or 2048×2048, with Z step sizes ranging from 1-3 μm. Most samples required multiple overlapping image stacks, which were then fused using ImageJ’s grid collection stitching plugin using the ‘unknown positions’ option [42]. Image stack downsampling, 3D segmentation and reconstruction were performed using the software Amira 6.1. For 3D reconstructions, pixels of each image stack were assigned to individual brain structures in the segmentation editor. We used the grey-value information of the anti-synapsin image stacks to define the boundaries of the neuropils in all three dimensions. The resulting voxel skeleton was then interpolated using Amira’s ‘wrap’ function, before generating a polygonal surface model to visualise the optic lobe in 3D. To improve clarity, all final images were adjusted for brightness and contrast in Adobe Photoshop CS6.

#### Layering analysis

We defined layers using confocal image stacks of a horizontal section through the optic lobe of *Hemicordulia tau*, co-labelled for synapsin and serotonin. A 45 μm z-stack was segmented into 6 maximum intensity projections of equal thickness, separating synapsin and serotonin channels. Two regions of interest were selected, one over the medulla and the other over the primary lobula. Care was taken to place these regions of interest in a position that minimised any shifts in structures across the X and Y planes at different depths. We used the plot profile tool in ImageJ to generate intensity plots for each region of interest, across both channels and at all 6 depths. Layers were segmented manually, defined by consistent changes in either synapsin or serotonin intensity across multiple depths of the sample.

## Author contributions

Conceptualization, J.M.F, D.C.O and B.e.J; Methodology, J.M.F, D.C.O and B.e.J.; Investigation, J.M.F, D.C.O and B.e.J.; Writing – Original Draft, J.M.F.; Writing – Review & Editing, J.M.F, D.C.O, S.D.W. and B.e.J.; Funding Acquisition, D.C.O and S.D.W.; Resources, D.C.O, S.D.W. and B.e.J.; Supervision, S.D.W. and D.C.O.

## Acknowledgements

This research was supported financially by the Swedish Research Council (VR 2014-4904, VR ‘-03452), the Australian Research Council (FF180100466, DP130104572) and STINT, the Swedish Foundation for International Cooperation in Research and Higher Education (STINT 2012-2033). We thank the manager of the Adelaide Botanic Gardens for allowing insect collection, and staff of Adelaide Microscopy and the Lund University Biology Microscopy facility for confocal microscope support.

## References

1. Misof, B, Liu, S, Meusemann, K, Peters, R.S., Donath, A, Mayer, C., Frandsen, P.B., Ware, J., Flouri, T., Beute, R.G. et al. (2014). Phylogenomics resolves the timing and pattern of insect evolution. Science. 346, 763–767.

2. Letch, H., Gottsberger, B., and Ware, J. L. (2016). Not going with the flow: a comprehensive time-calibrated phylogeny of dragonflies (Anisoptera: Odonata: Insecta) provides evidence for the role of lentic habitats on diversification. Molecular Ecology. 25, 1340–1353.

3. Zawarzin, A. (1914). Histologische Studien über Insekten. IV. Die optischen Ganglien der Aeschna-Larven. Z. Wiss. Zool. 108: 175–257

4. Cajal, S. R., and Sanchez D (1915) Contribucion al conocimiento de los centros nerviosos de los insectes. Trab Lab Invest Biol Univ Madrid. 13, 1–168.

5. Strausfeld, N. (2009). Brain organization and the origin of insects: an assessment. Proc. R. Soc. B. 276, 1929–1937.

6. Ito, K., Shinomiya, K., Ito, M., Armstrong, J. D., Boyan, G., Hartenstein, V., Harzsch, S., Heisenberg, M., Homberg, U., Jenett, A., Keshishian, H., Restifo, L. L., Rössler, W., Simpson, J. H., Strauss, R., Vosshall, L. B., et al. (2014). Insect Brain Name Working Group. A systematic nomenclature for the insect brain. Neuron. 19, 755–65.

7. Strausfeld, N. (2005). The evolution of crustacean and insect optic lobes and the origins of chiasmata. Arthropod Struct & Dev. 34, 235–256.

8. Klagges, B. R., Heimbeck, G., Godenschwege, T. A., Hofbauer, A., Pflugfelder, G. O., Reifegerste, R., Reisch, D., Schaupp, M., Buchner, S., and Buchner, E. (1996). Invertebrate synapsins: a single gene codes for several isoforms in *Drosophila*. J Neurosci 16, 3154–3165.

9. Pritchard, G. (1966). On the morphology of the compound eyes of dragonflies (Odonata: Anisoptera), with special reference to their role in prey capture. Physiological Entomology. 41, 1–8.

10. Rein, K., Zöckler, M., Mader, M.T., Grübel, C., Heisenberg, M. (2002). The drosophila standard brain. Current Biol. 3, 227–231.

11. Immonen, E., Dacke, M., Heinze, S., el Jundi, B. (2017). Anatomical organization of the brain of a diurnal and a nocturnal dung beetle. J Comp Neurol. 525, 1879–1908.

12. Dreyer, D., Vitt, H., Dippel, S., Goetz, B., el Jundi, B., Kolimann, M., Huetteroth, W., Schachtner, J. (2010). 3D standard brain of the red flour beetle *Tribolium castaneum*: a tool to study metamorphic development and adult plasticity. Front Syst Neurosci. 4, 3.

13. Kvello, P., Løfaldli, B., Rybak, J., Menzel, R., and Mustaparta, H. (2009). Digital, three-dimensional average shaped atlas of the *Heliothis virescens* brain with integrated gustatory and olfactory neurons. Front Syst Neurosci 3, 14.

14. el Jundi, B., Huetteroth, W., Kurylas, A.E., and Schachtner J. (2009). Anisometric brain dimorphism revisited: implementation of a volumetric 3D standard brain in *Manduca sexta*. J Comp Neurol, 517, 210–225.

15. Heinze, S, and Reppert SM. (2012). Anatomical basis of sun compass navigation I: the general layout of the monarch butterfly brain. J Comp Neurol. 520, 1599–1628.

16. Montgomery, S. H., and Ott, S R. (2015). Brain Composition in *Godyris zavaleta*, a Diurnal Butterfly, Reflects an Increased Reliance on Olfactory Information. J Comp Neurol. 523, 869–891.

17. Stöckl, A., Heinze, S., Charalabidis, A., el Jundi, B., Warrant, E., and Kelber, A. (2016). Differential investment in visual and olfactory brain areas reflects behavioural choices in hawk moths. Sci Rep. 6, 26041.

18. Fischbach KF, Dittrich APM. (1989). The optic lobe of *Drosophila melanogaster*. I: A. Golgi analysis of wild-type structure. Cell Tissue Res. 258, 441–475.

19. Ribi, W. A., and Scheel, M. (1981). The second and third optic ganglia of the worker bee. Cell Tissue Res. 221, 17–43.

20. Brandt, R., Rohlfing, T., Rybak, J., Krofczik, S., Maye, A., Westerhoff, M., Hege, H., Menzel, H. (2005). Three-Dimensional Average-Shape Atlas of the Honeybee Brain and Its Applications. J Comp Neurol. 492, 1–19.

21. Kinoshita, M., Shimohigasshi, M., Tominaga, T., Arikawa, K., and Homberg, U. (2015). Topographically Distinct Visual and Olfactory Inputs to the Mushroom Body in the Swallowtail Butterfly, *Papilio xuthus*. J. Comp. Neurol. 523, 162–182.

22. Rosner, R., von Haldeln, J., Salden, T., and Homberg, U. (2017). Anatomy of the lobula complex in the brain of the praying mantis compared to the lobula complexes of the locust and cockroach. J. Comp. Neurol. 525, 2343–2357.

23. Strausfeld, N. (1970). Golgi studies on insects. Part II. The optic lobes of diptera. Philos Trans R Soc Lond B Biol Sci. 258, 135–223.

24. Nässel, D. R. and Klemm, N. K. (1983). Serotonin-like immunoreactivity in the optic lobes of three insect species. Cell Tissue Res. 232, 129:140.

25. Wendt, B, and Homberg, U. (1992). Immunocytochemistry of dopamine in the brain of the locust Schistocerca gregaria. J Comp Neurol. 321, 387–403.

26. Geurten, B.R.H., Nordström, K., Sprayberry, J.D.H., Bolzon, D.M., and O’Carroll, D.C. (2007). Neural mechanisms underlying target detection in a dragonfly centrifugal neuron. J. Exp. Biol. 210, 3277–3284.

27. Wiederman, S. D., Fabian, J.M., Dunbier, J.R., & O’Carroll, D.C. (2017) A predictive focus of gain modulation encodes target trajectories in insect vision eLife, doi: 10.7554/eLife.26478 (in press)

28. Gilbert, C., and Strausfeld, N. J. (1991). The functional organization of male-specific visual neurons in flies. J Comp Physiol A. 169, 395–411.

29. Maddess, T., and Yang, E. (1997). Orientation-sensitive Neurons in the Brain of the Honey Bee (Apis mellifera). J Insect Physiol. 43, 329–336.

30. Nordström, K., and O’Carroll, D. C. (2009). Feature detection and the hypercomplex property in insects. Trends Neurosci. 32, 383–391.

31. El Jundi, B., Pfeiffer, Keram., Heinze, and S., Homberg, U. (2014). Integration of polarization and chromatic cues in the insect sky compass. J Comp Physiol A. 200, 575–589.

32. Hausen, K. (1982). Montion Sensitive Interneurons in the Optopmotor System of the Fly. I. The Horizontal Cells: Structure and Signals. Biol. Cybern. 45, 143–156.

33. Evans, B. J. E., O’Carroll, D. C., Fabian, J. M., and Wiederman, S. D. (2019). Differential Tuning to Visual Motion Allows Robust Encoding of Optic Flow in the Dragonfly. J. Neurosci. 39, 8051–8063.

34. DeVoe, R. D., Kaiser, W., Ohm, J., and Stone, L. S. (1982). Horizontal movement detectors of honeybees: Directionally-selective visual neurons in the lobula and brain. J. Comp Physiol. 147, 155–170.

35. Ibbotson, M. R. (1991) Wide-field motion-sensitive neurons tuned to horizontal movement in the honeybee, *Apis mellifera*. J. Comp. Physiol. A. 168, 91–102.

36. Shinomiya, K., Takemura, Shin-ya., Rivlin, P. K., Plaza, S. M., Scheffer, L. K., and Meinertzhagen, I A. (2015). A common evolutionary origin for the ON-and OFF-edge motion detection pathways of the Drosophila visual system. Front. Neural Circuits. 9, 33.

37. Ott, S. R. (2008). Confocal microscopy in large insect brains: Zinc-formaldehyde fixation improves synapsin immunostaining and preservation of morphology in whole-mounts. J Neurosci Methods. 172, 220–230.

38. Gonzalez-Bellido, P. T., Wardill, T. J. (2012). Labeling and Confocal Imaging of Neurons in Thick Invertebrate Tissue Samples. Cold Spring Harb Protoc.

39. Wiederman, S. D., Fabian, J.M., Dunbier, J.R., & O’Carroll, D.C. (2017) A predictive focus of gain modulation encodes target trajectories in insect vision eLife, doi: 10.7554/eLife.26478

40. Fabian, J.M., Dunbier, J.R., O’Carroll, D.C., & Wiederman, S. D. (2019) Properties of predictive gain modulation in a dragonfly visual neuron. J. Exp. Biol. 222, jeb207316.

41. Lancer, B.H., Evans, B.J.E., Fabian, J.M., O’Carroll, D.C., & Wiederman, S.D. (2019) A Target-Detecting Visual Neuron in the Dragonfly Locks on to Selectively Attended Targets. J. Neurosci. 39, 8497–8509.

42. Preibisch, S., Saalfeld, S., & Tomancak, P. (2009) Globally optimal stitching of tiled 3D microscopic image acquisitions. Bioinformatics (2009) 25: 1463–1465. DOI: https://doi.org/10.1093/bioinformatics/btp184

