## Supplemental Information for "The complex optic lobe of dragonflies"

Figure S1. Related to Figure 1

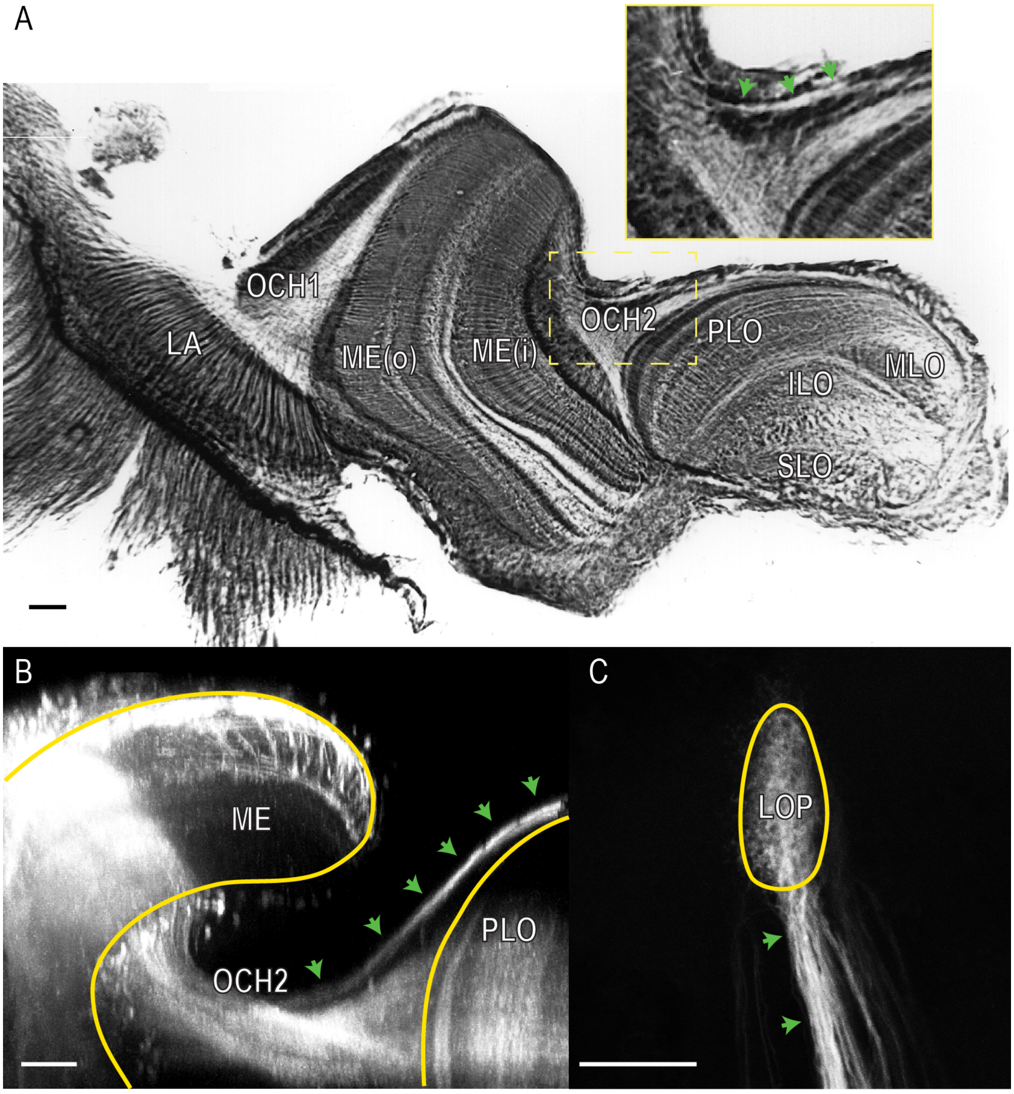


A) A horizontal section through the optic lobe of *H. tau*, stained with osmium tetroxide. Osmium binds to lipids in cellular membranes, resulting in dark staining in membrane rich areas. Unlike synapsin immunolabelling, this technique allows the visualisation of both synaptic neuropil and axons, allowing the identification of major axonal tracts connecting brain regions, whilst still allowing recognition of synaptically dense regions (darker staining) in neuropils identified using anti-synapsin labelling (Figure 1). Lamina output axons enter the medulla via the first optic chiasm (OCH1). Medulla outputs enter the lobula complex by two separate routes: most axons enter the primary lobula via the second optic chiasm (OCH2), whilst a smaller number of axons take a more posterior route, running across the posterior surface of the primary lobula (inset, green arrows). The lobula plate is not visible at the level of this section. B) An XZ view (Y projection) through the optic lobe of *A. mixta* from a confocal image stack (Z series) obtained from a whole-mount brain following a dextran tracer injection into the posterior medulla. Fluorescence reveals a large number of columnar neurons projecting from the inner medulla and then entering the outer primary lobula (PLO) via the second optic chiasm (OCH2) and a second minor tract projecting across and bypassing the posterior part of the primary lobula (green arrows). C) An XY view (Z projection) of the posterior processes from the same Z series, showing that they terminate in the lobula plate (LOP), on the posterior surface of the lobula complex. Scale bars = 50µm.

Table S1. Related to Figure 2

| | *H. tau* | | *S. striolatum* | | *A. mixta* | |
|---|---|---|---|---|---|---|
| Structure | Absolute volume (μm$^3$) | Relative Volume (% total) | Absolute volume (μm$^3$) | Relative Volume (% total) | Absolute volume (μm$^3$) | Relative Volume (% total) |
| Lobula Plate | $2.92 \times 10^6$ | 0.38 | $1.69 \times 10^6$ | 0.38 | $3.04 \times 10^6$ | 0.34 |
| Lobula (other) | $9.91 \times 10^7$ | 14.54 | $5.99 \times 10^7$ | 13.60 | $1.14 \times 10^8$ | 12.95 |
| Remaining brain (excluding lamina) | $5.80 \times 10^8$ | 85.08 | $3.79 \times 10^8$ | 86.02 | $7.66 \times 10^8$ | 86.71 |
| **Total** | **$6.81 \times 10^8$** | **100** | **$4.40 \times 10^8$** | **100** | **$8.83 \times 10^8$** | **100** |

Figure  S2. Related to Figure 3

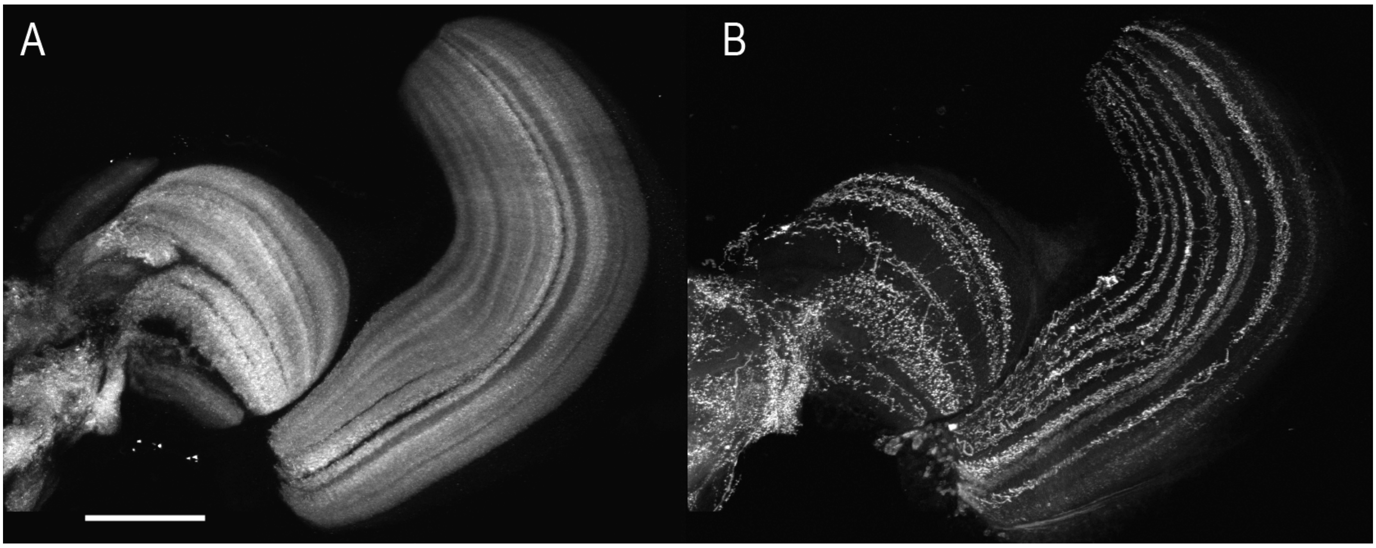


Channels separated from a maximum intensity (Z) projection of a stack of confocal images obtained from stained horizontal section through the optic lobe in *H. tau* as used for the layering analysis (Figure 3). A) the synapsin labelled channel. B) The serotonin channel from the same section. Scale bars = 200µm.
